# Activity-dependent LTP in the dentate gyrus promotes epileptic seizures

**DOI:** 10.1101/2021.06.30.450628

**Authors:** Kaoutsar Nasrallah, M. Agustina Frechou, Young J. Yoon, Subrina Persaud, Tiago Gonçalves, Pablo E. Castillo

## Abstract

Epilepsy is a devastating brain disorder whose cellular mechanisms remain poorly understood. Excitatory mossy cells (MCs) in the dentate gyrus of the hippocampus are implicated in temporal lobe epilepsy, the most common form of focal epilepsy in adults. However, the role of MCs during initial seizures, before MC loss occurs, is unclear. Here, we show that initial seizures induced with kainic acid (KA) intraperitoneal injection in adult mice, a well-established model of experimental epilepsy, not only increased MC and granule cell (GC) activity *in vivo*, but also triggered a BDNF-dependent long-term potentiation at MC-GC synapses (MC-GC LTP). *In vivo* induction of MC-GC LTP worsened KA-induced seizures, whereas selective MC silencing and *Bdnf* genetic removal from GCs, which abolishes LTP, were both anti-epileptic. Thus, initial seizures strengthen MC-GC synaptic transmission, thereby promoting epileptic activity. Our findings reveal a potential mechanism of epileptogenesis that may help develop therapeutic strategies for early intervention.

## INTRODUCTION

Epilepsy is a common neurological disorder characterized by recurrent epileptic seizures, often associated with profound cognitive, psychological, and social deleterious consequences^1^. About 30 % of the patients are resistant to antiepileptic drugs that do not necessarily target pathogenic mechanisms involved in epileptogenesis^2^. To develop more effective treatments, a better understanding of the cellular and molecular processes implicated in the early stages of epilepsy, before the brain damages become irreversible, is required. Mossy cells (MCs), excitatory neurons in the dentate gyrus (DG) of the hippocampus, play a critical role in temporal lobe epilepsy (TLE)^3–5^, the most common form of focal epilepsy in adults^6^. However, whether MC activity can be pro- or anti-epileptic has been a subject of debate for several decades^3–5,7–10^. MC loss is a hallmark feature of chronic TLE in both humans and animal models^11–15^. Recent studies reported that while surviving MCs in a mouse model of chronic TLE play an antiepileptic role^4^, these cells could be pro-epileptic early during initial experimental seizures^5^.

Recurrent excitatory activity is a core mechanism in epilepsy^16^. In the DG, glutamatergic MCs and granule cells (GCs) are reciprocally connected thus forming an intrinsic excitatory loop. Remarkably, a single MC makes more than 30,000 synaptic contacts onto GCs, locally, contralaterally, and along the longitudinal axis of the hippocampus^17,18^. Furthermore, repetitive stimulation of MC axons *in vitro*, induces robust long-term potentiation at MC-GC excitatory synapses (MC-GC LTP). The DG is characterized by very sparse activation of GCs^19–21^ and it is believed to act as a gate that opens during epileptic seizures^22,23^. The long-lasting strengthening of MC-GC synaptic transmission is sufficient to overcome the basal strong GC inhibition, thereby allowing MCs to drive GCs and presumably open the DG gate^24^. MC-GC LTP is mediated by brain-derived neurotrophic factor (BDNF)/Tropomyosin receptor kinase B (TrkB) signaling^24,25^, which is known to promote TLE^26,27^. Therefore, activity-dependent strengthening of MC-GC synapses may promote epilepsy through the extensive MC projections onto GCs. While an episode of prolonged seizures (e.g. *status epilepticus*) can result in TLE^28–31^, it is unknown whether and how initial seizures can impact MC-GC synaptic strength.

Using multiple complementary approaches, such as chemogenetics, *in vitro* electrophysiology, *in vivo* optogenetics, *in vivo* calcium imaging and a conditional knockout strategy, we found that initial seizures not only increased MC and GC activity, *in vivo*, but also triggered a BDNF-dependent strengthening of MC-GC synaptic transmission. In addition, *in vivo* induction of MC-GC LTP was sufficient to promote convulsive seizures, whereas interfering with BDNF signaling and MC activity had an anti-convulsant effect. Our findings support a pro-epileptic role of MCs and BDNF in early TLE and provide a potential causal link between MC-GC LTP and epilepsy.

## RESULTS

### Chemogenetic silencing of MCs reduced acute kainic acid-induced seizures

To test the hypothesis that MC-GC LTP has a pro-convulsant effect during early epilepsy, we first tested the prediction that silencing MCs should reduce the severity/susceptibility of experimental seizures induced by a single intraperitoneal (IP) injection of kainic acid (KA, 30 mg/kg), a well-established experimental model of epilepsy^32,33^. To suppress MC output the G_i_ (hM4Di) inhibitory designer receptor exclusively activated by designer drug (iDREADD) was selectively expressed in MCs. We bilaterally injected a Cre-recombinase-dependent virus expressing the iDREADD (AAV-CaMKII-DIO-hM4D(G_i_)-mCherry) into the DG of *Drd2*-Cre mice, whereas *Drd2*-Cre mice injected with AAV-CaMKII-DIO-mCherry served as control (Fig 1A). Consistent with previous reports^5,34^, we found that the viral expression was selective to MCs (Fig 1B). Next, we verified that the iDREADD efficiently responded to the DREADD agonist clozapine N-oxide (CNO). Bath application of 10 μM CNO significantly reduced MC-GC EPSC amplitude in acute slices obtained from *Drd2*-Cre mice injected with AAV-CaMKII-DIO-hM4D(G_i_)-mCherry but not with AAV-CaMKII-DIO-mCherry (Fig 1 C and D, iDREADD: 43.8 ± 4.3 % of baseline, n = 6, p < 0.001, paired t-test; control: 100.6 ± 5.5 % of baseline, n = 5, p > 0.05, paired t-test). We then monitored and scored behavioral seizures induced by KA IP injection (30 mg/kg) for 2 hours (see Methods) in both *Drd2*-Cre mice injected with AAV-CaMKII-DIO-hM4D(G_i_)-mCherry (iDREADD) or AAV-CaMKII-DIO-mCherry (control). The two groups were injected with CNO (2 mg/kg) 30 minutes prior KA IP administration (30 mg/kg). Consistent with a recent study using the pilocarpine-model^5^, we found that silencing MCs reduced seizure severity and susceptibility, as indicated by significant decrease in the total cumulative seizure score (Fig 1 F, two way ANOVA RM, AAV condition: F(1, 5) = 7.2, p < 0.05, time: F(1.1, 5.6) = 71.7, p < 0.001; AAV condition x time: F(1,5) = 88.1, p < 0.001), in sum score (Fig 1 G, control: 33.7 ± 5.1, n = 7; iDREADD: 17.8 ± 3.2, n = 6; control *vs* iDREADD: p < 0.05, unpaired t-test) and increase in latency to stage 3 (Fig 1 H, control: 45.7 ± 6.1, n = 7; iDREADD: 91.7 ± 14.9, n = 6; control *vs* iDREADD: p < 0.05, unpaired t-test). These results reinforce the notion that MC activity has a pro-convulsant effect during initial epileptic seizures.

**Figure 1:**
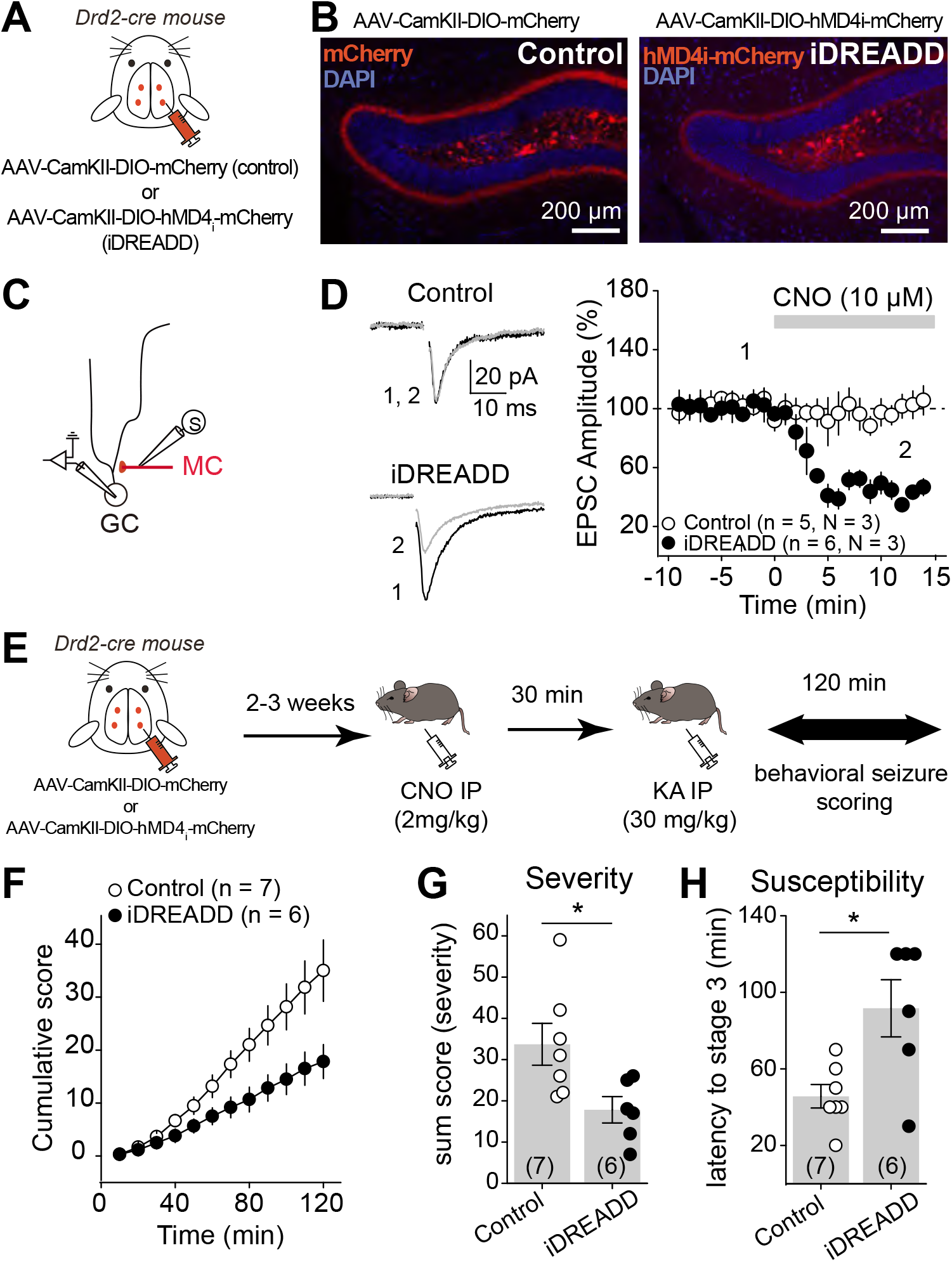
Chemogenetic silencing of MCs reduced acute kain acid-induced seizures. **(A, B)** AAV expressing mCherry (control, AAV-CaMKII-DIO-mCherry) or the inhibitory DREADD hMD4_i_-mCherry (iDREADD, AAV-CaMKII-DIO-hMD4_i_-mCherry) were injected bilaterally into the ventral and dorsal DG of Drd2-cre mice. Confocal images of the DG showing how the viral expression is selective for hilar MCs. Note the dense labeling of MC axons in the inner molecular layer (IML). **(C, D)** Schematic diagram illustrating the recording configuration (C). MC EPSCs were recorded from GC in response to MC axon stimulation in the IML. (D) Representative traces (left) and time course plot (right) showing that CNO significantly reduced EPSC amplitude in slices expressing the inhibitory DREADD in MCs but not in controls. Numbers in parentheses indicate the number of cells (n) and the number of animals (N). **(E)** Representation of the experimental timeline. Viral stereotaxic injections were performed in Drd2-cre mice to express control (AAV-CaMKII-DIO-mCherry) or iDREADD in MCs (AAV-CaMKII-DIO-hMD4_i_-mCherry) 2-3 weeks before assessing behavioral seizures (for 120 min). All animals were treated with CNO *in vivo* (2 mg/kg, IP) 30 min before seizures were acutely induced with single KA IP injection (30 mg/kg). **(F-H)** Chemogenetic silencing of MCs reduced seizure severity and susceptibility. Scoring of seizures using a modified Racine scale for 120 min revealed a significant decrease in cumulative seizure score (F), in sum score (G) and a significantly increase in latency to convulsive seizures (H) when MCs were silenced as compared with control animals. Numbers in parentheses indicate the number of animals. * p < 0.5. Data are presented as mean ± SEM.

### Initial convulsive seizures potentiated MC-GC transmission presynaptically

We hypothesized that initial epileptic seizures may increase MC activity, and thus induce MC-GC LTP *in vivo*. MC repetitive activity is sufficient to trigger a robust MC-GC LTP in acute brain slices obtained from healthy rodents^24^. Furthermore, a recent *in vivo* study using calcium indicator and fiber photometry reported that DG neuronal activity is increased during KA-induced seizures^35^. However, the contribution of specific subtypes of neurons, including MCs and GCs, remains unknown. We therefore monitored MC and GC activity *in vivo* using calcium imaging during acutely induced seizures. To this end, we expressed the genetically encoded Ca^2+^ indicator jRGECO1a selectively in DG excitatory neurons, by unilaterally injecting AAV-CaMKII-jRGECO1a into the DG of wild type (WT) adult mice. The animals were then implanted with a chronic imaging window above the dorsal hippocampus and MC and GC activity was visualized using head-fixed 2 photon imaging before and during acute seizures (Fig 2A, B). After collecting basal activity, seizures were induced with a single KA IP injection (20 mg/kg). Neuronal activity was monitored during stage 3 of convulsive seizures, which was determined by the presence of forelimb clonus. Saline-injected mice served as a control. In total, we recorded 94 MCs and 136 GCs from 4 saline-injected mice and 63 MCs and 66 GCs from 5 KA-injected mice. We found that both MC (Fig 2 C and D, saline: −0.06 ± 0.07, N = 4, p > 0.05, t-test; KA: 7.6 ± 1.4, N = 5, p < 0.01, t-test) and GC (Fig 2 C and D, saline: −0.04 ± 0.08, N = 4, p > 0.05, t-test; KA: 6.4 ± 1.8, N = 5, p < 0.05, t-test) calcium signals were significantly increased during KA-induced convulsive seizures but not after saline administration, indicating a robust increase in neuronal activity.

**Figure 2:**
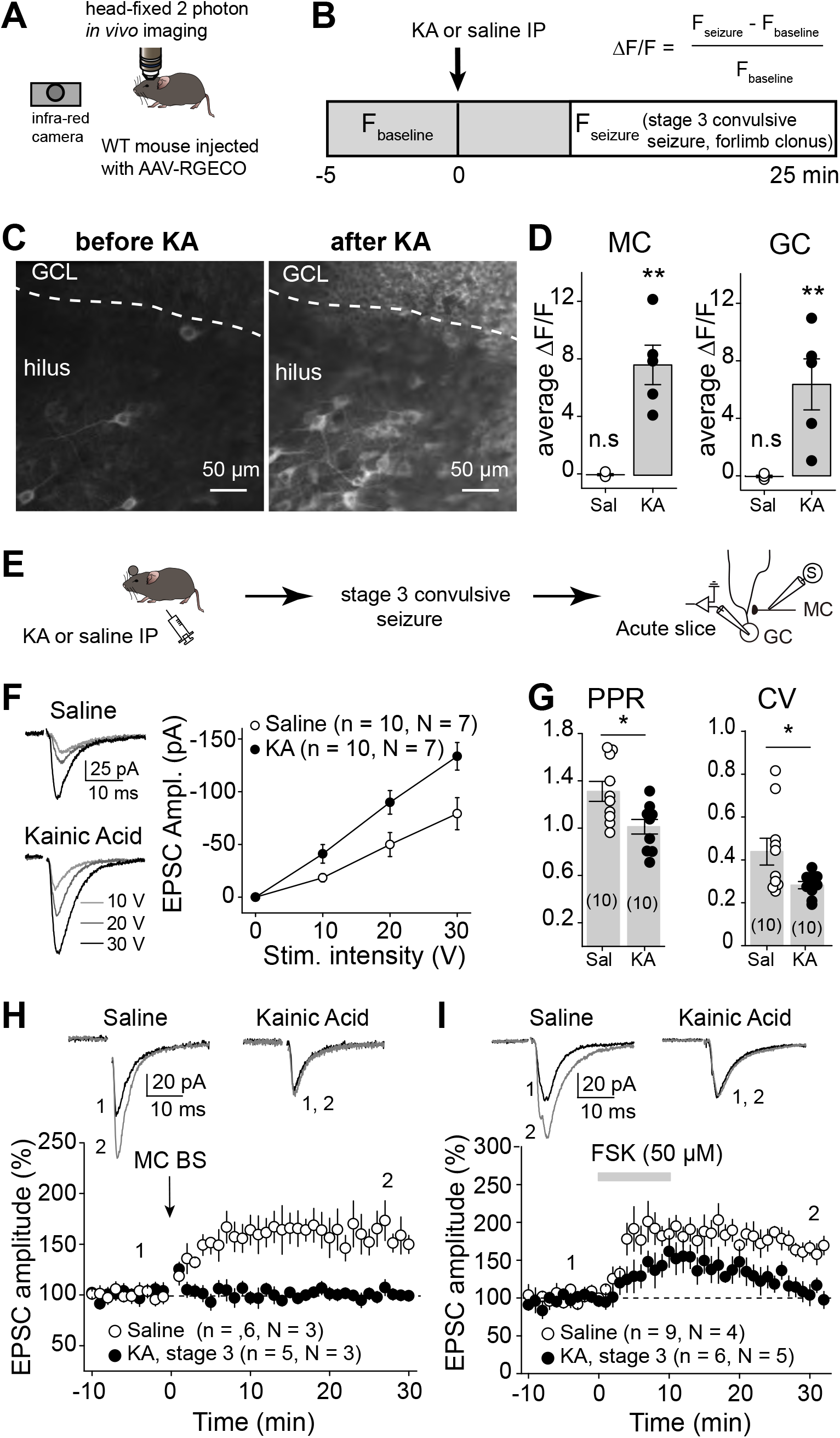
Initial convulsive seizures increased DG excitatory neuron activity and MC-GC synaptic strength. **(A, B)** Schematic diagram showing the experimental apparatus (A). jRGECO1a-expressing MCs and GCs were imaged before and during KA (30 mg/kg)-induced convulsive seizures, in head-fixed mice monitored with IR camera. Saline injections were used as control. **(C, D)** Mean Z-projection image, acquired *in vivo*, of jRGECO1a-expressing MCs (hilus) and GCs (granule cell layer, GCL), before and after KA injection (C). Average ΔF/F of recorded MCs and GCs after saline or KA injection (D). Each data point corresponds to the average value per animal. **(E)** Seizures were acutely induced using KA IP (20 mg/kg). The mice were sacrificed once stage 3 of convulsive seizures was reached and MC-GC synaptic function was accessed in acute hippocampal slices. Saline-injected mice (IP) were used as control. **(F)** Representative traces and summary plot showing how input/output function was increased in KA (20 mg/kg)-injected mice. Numbers in parentheses represent number of cells (n) and the number of mice (N). **(G)** PPR and CV were both significantly decreased in KA-treated mice as compared to saline-injected mice. Numbers in parentheses represent number of cells. **(H, I)** Representative traces (left) and time-course summary plots (right) showing that LTP at MC-GC synapse induced by either MC BS (5 pulses at 100Hz, repeated 50 times every 0.5 s, H) or 50 μM forskolin (FSK, I) application was impaired in KA-injected mice (20 mg/kg KA, stage 3). Numbers in parentheses indicate the number of cells (n) and the number of mice (N). * p < 0.5, ** p < 0.01. Data are presented as mean ± SEM.

We then tested whether initial epileptic seizures, by increasing MC activity and inducing LTP, could have strengthen MC-GC connections *in vivo*. We analyzed MC-GC synaptic transmission in both KA-injected and saline-injected mice. After a single KA injection (20 mg/kg, IP), the animals were monitored and sacrificed for acute hippocampal slice preparation once stage 3 of convulsive seizures was reached (Fig 2E). Sham-injected mice were used as control. MC-GC synaptic function was assessed by activating MC axons while performing whole-cell voltage-clamp recordings from GCs. We found that MC-GC synaptic transmission was significantly strengthened, as indicated by an increase in the input/output function (Fig 2F, two way ANOVA RM, IP injection: F(1, 9) = 6.5, p < 0.05, stimulation intensity: F(1.4, 12.5) = 64.5, p < 0.001; IP injection x stimulation intensity: F(1, 9) = 139.8, p < 0.001), while both paired-pulse ratio (PPR) and coefficient of variation (CV) were significantly reduced in KA-injected as compared to saline-treated mice (Fig 2G, PPR: saline: 1.31 ± 0.08, n = 10; KA: 1.01 ± 0.06, n = 10; saline *vs* KA: p < 0.05, unpaired t-test; CV: saline: 0.44 ± 0.06, n = 10; KA: 0.28 ± 0.02, n = 10; saline *vs* KA: p < 0.05, unpaired t-test). These results strongly suggest that initial seizures, presumably by inducing presynaptic LTP, strengthened MC-GC synapses *in vivo*. If so, this plasticity should be occluded in hippocampal slices prepared from KA-injected mice. In support of this hypothesis, we found that both synaptically-induced LTP by repetitive stimulation of MC axons (Fig 2H, saline: 155.5 ± 14.7 % of baseline, n = 6, p < 0.05, paired t-test; KA: 99.2 ± 3.8 % of baseline, n = 5, p > 0.05, paired t-test; saline *vs* KA: p < 0.05, unpaired t-test) and chemically-induced LTP by transient application (10 min, 50 μM) of the adenylyl-cyclase activator forskolin^24^ (Fig 2I, saline: 170.7 ± 7.6 % of baseline, n = 9, p < 0.001, paired t-test; KA: 114.3 ± 9.8 % of baseline, n = 6, p > 0.05, paired t-test; saline *vs* KA: p < 0.001, unpaired t-test), were impaired in KA-injected mice as compared to control (Fig 2H and I). Altogether, our findings strongly suggest that early KA-induced seizures strengthened MC-GC synaptic transmission by inducing MC-GC LTP *in vivo*.

### Blocking seizure-induced MC-GC LTP had an anti-convulsant effect

BDNF/TrkB signaling is critical for MC-GC LTP. BDNF is released, by both MCs and GCs, upon repetitive presynaptic activity and is necessary and sufficient for the induction of MC-GC LTP (Berthoux et al.)^24^. To test whether BDNF is also involved in the seizure-induced strengthening of MC-GC synaptic transmission, occurring *in vivo*, we conditionally KO *Bdnf* from GC, a manipulation that abolishes MC-GC LTP (Berthoux et al.), and tested whether seizures can trigger MC-GC potentiation in absence of postsynaptic BDNF. Given that experimental seizures can be reduced in *Bdnf* KO mice^36,37^, we only injected 0.5 μl of AAV5.CaMKII.Cre-mCherry or AAV5.CaMKII.mCherry (control) into the DG upper blade of *Bdnf*^fl/fl^ mice in order to prevent a potential failure in seizure induction when knocking out *Bdnf* from DG excitatory neurons. Mice were sacrificed 25 min after KA injection, which is the average time for reaching stage 3 convulsive seizure in WT control mice. We then prepared acute slice to monitor MC-GC synaptic function (Fig 3A). While behavioral seizures were comparable in control and cKO mice (all animals reached stage 3 of convulsive seizures by minute 25), *Bdnf* deletion from GCs (Cre-mCherry+ GCs) prevented seizure-induced MC-GC LTP, as KA IP injection failed to increase MC EPSC amplitude (Fig 3B, two way ANOVA RM, IP injection: F(1, 6) = 0.003, p > 0.05, stimulation intensity: F(1.56, 9.38) = 57.97, p < 0.001; IP injection x stimulation intensity: F(1,6) = 287.4, p < 0.001) or decrease PPR (Fig 3C, saline: 1.2 ± 0.01, n = 7; KA: 1.2 ± 0.1, n = 7; saline vs KA: p > 0.05, unpaired t-test) and CV (Fig 3C, saline: 0.40 ± 0.04, n = 7; KA: 0.42 ± 0.06, n = 7; saline *vs* KA: p > 0.05, unpaired t-test) as compared to saline-injected mice. The lack of KA-induced synaptic strengthening was not due to viral expression since KA injection efficiently increased the input/output function (MC EPSC amplitude, Fig 3D, two way ANOVA RM, IP injection: F(1, 4) = 23.6, p < 0.01, stimulation intensity: F(1.34, 5.34) = 68.9, p < 0.001; IP injection x stimulation intensity: F(1,4) = 537.0, p < 0.001) and reduced PPR (Fig 3E, saline: 1.34 ± 0.1, n = 5; KA: 0.98 ± 0.1, n = 7; saline *vs* KA: p > 0.05, unpaired t-test) and CV (Fig 3E, saline: 0.39 ± 0.04, n = 5; KA: 0.27 ± 0.03, n = 7; saline *vs* KA: p < 0.05, unpaired t-test) in mCherry+ control GCs. Of note, input/output function, PPR and CV in cKO (Cre-mCherry+) GCs and control (mCherry+) GCs after saline injection were comparable (Fig 3B-E). GC membrane properties were similar between all the different groups (Fig S1). These data suggest that while *Bdnf* cKO had no impact on basal MC-GC synaptic properties and GC membrane properties, it abolished seizure-induced strengthening of MC-GC synaptic transmission.

**Figure 3:**
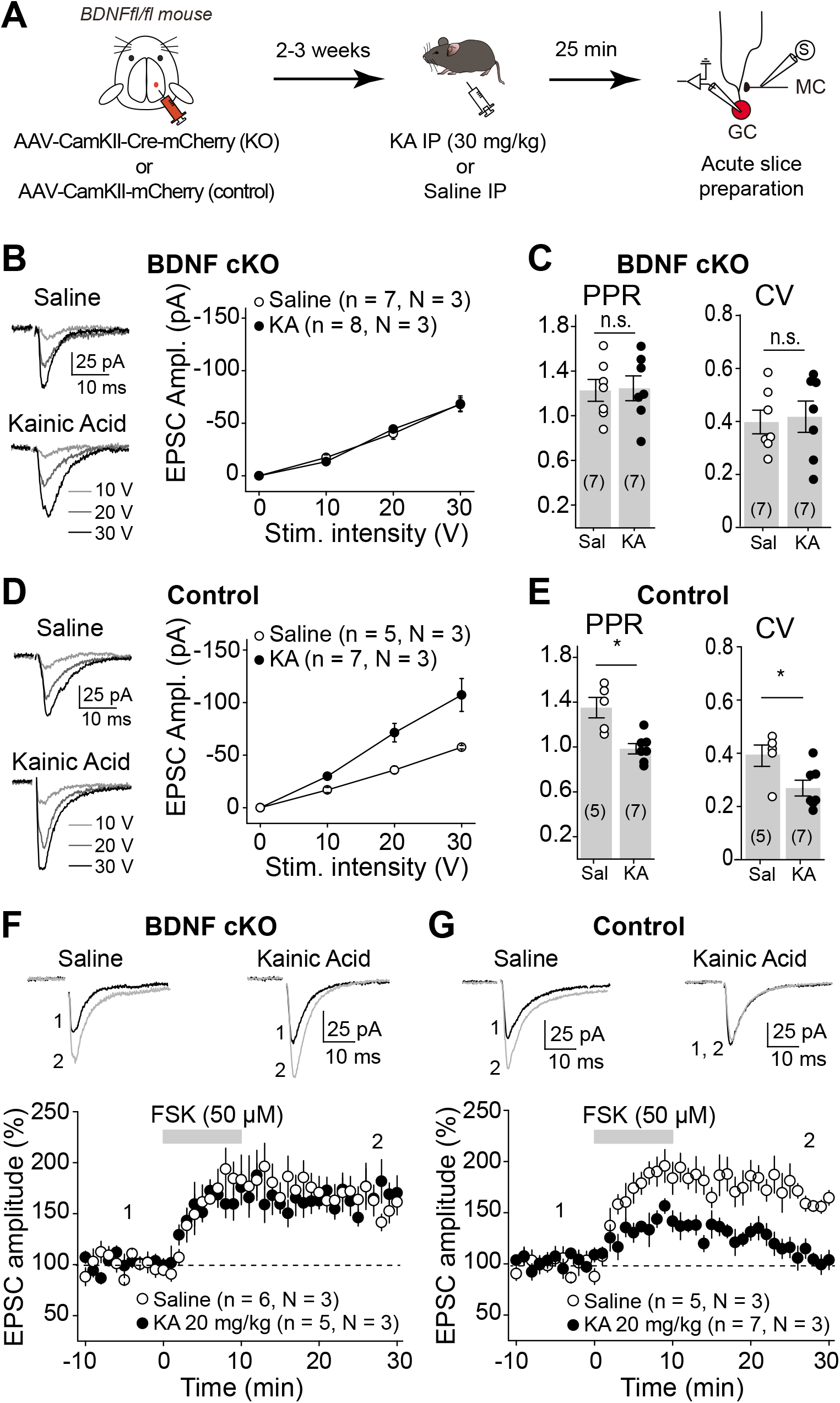
Seizure-induced MC-GC synaptic strengthening required postsynaptic BDNF. **(A)** Representation of the experimental timeline. Control (AAV-CaMKII-mCherry) or Cre-expressing AAV (AAV-CaMKII-Cre-mCherry) were injected unilaterally into the dorsal blade of the DG of *Bdnf*^fl/fl^ mice. Seizures were induced 2-3 weeks later using KA IP (30 mg/kg) and acute hippocampal slices were prepared 25 min post-injection. We then performed whole cell recordings in mCherry+ GC, in response to MC axon stimulation. **(B, C)** KA-induced seizure failed to increase MC-GC synaptic strength in GC lacking BDNF. EPSC Amplitude (B), PPR (C) and CV (C) were similar in Cre-mCherry+ GC obtained from KA vs saline-injected mice. **(D, E)** Seizure induction triggered a normal increase in MC-GC EPSC amplitude (E) and decrease in PPR (E) and CV (E) in *Bdnf*^fl/fl^ mice injected with a control virus (AAV-CaMKII-mCherry). **(F)** Representative traces and time-course summary plots showing that bath application of forskolin (FSK, 50 μM, 10 min) induced similar LTP in KA- and saline injected mice when BDNF was KO from GC (*Bdnf*^fl/fl^ mice injected with AAV-CaMKII-Cre-mCherry). **(G)** LTP at MC-GC synapse induced by forskolin (FSK) was impaired in KA-injected mice (20 mg/kg KA) as compared to saline-treated mice, in *Bdnf*^fl/fl^ mice injected with a control virus (AAV-CaMKII-mCherry). In panels C and E, numbers in parentheses represent number of cells. In panels B, D, F and G, numbers in parentheses indicate the number of cells (n) and mice (N). * p < 0.05, n.s. p > 0.05. Data are presented as mean ± SEM.

Because PKA activity is required for MC-GC LTP downstream of BDNF/TrkB signaling^24^(Berthoux et al.,), we examined whether cAMP/PKA activation could still induce LTP in BDNF-deficient GCs. Bath application of forskolin (50 μM, 10 min) triggered normal LTP in BDNF-deficient GCs (Cre-mCherry+) obtained from both KA- and saline-injected mice in postsynaptic *Bdnf* cKOs (Fig 3F, saline: 162.9 ± 9.4 % of baseline, n = 6, p < 0.01, paired t-test; KA: 161.9 ± 5.7 % of baseline, n = 5, p < 0.001, paired t-test; saline *vs* KA: p > 0.05, unpaired t-test), supporting the idea that postsynaptic *Bdnf* deletion prevented KA-injection from inducing MC-GC LTP *in vivo*. In contrast, and consistent with our previous results (Fig 2I), forskolin failed to induce LTP in control GCs (mCherry+) obtained from KA-injected mice but not from saline-injected mice (Fig 3G, saline: 171.5 ± 7.2 % of baseline, n = 5, p < 0.001, paired t-test; KA: 115.7 ± 8.1 % of baseline, n = 7, p < 0.05, paired t-test; saline *vs* KA: p < 0.001, unpaired t-test). Altogether, these results reveal that initial seizures triggered MC-GC LTP *in vivo* via a BDNF-dependent mechanism, and that such synaptic potentiation occluded subsequent induction of LTP *in vitro*.

Deleting *Bdnf* from DG excitatory neurons, a manipulation that abolishes MC-GC LTP both *in vitro* (Berthoux et al) or *in vivo* (Fig 3), could inhibit seizure induction. To test this possibility, we stereotaxically injected AAV-CaMKII-Cre-mCherry (cKO) or AAV-CaMKII-mCherry (control) into both dorsal and ventral DG of adult *Bdnf*^fl/fl^ mice, bilaterally (Fig 4A). We confirmed that the virus was highly expressed in the DG (Fig 4B) and found that at least 70% of DG neurons were mCherry+. *Bdnf* conditional deletion reduced seizure severity and susceptibility, as indicated by significant decrease in the total cumulative seizure score (Fig 4C, two way ANOVA RM, AAV condition: F(1, 7) = 6.0, p < 0.05, time: F(1.26, 8.87) = 150.4, p < 0.001; AAV condition x time: F(1,7) = 262.4, p < 0.001), in sum score (Fig 4D, control: 48.9 ± 6.0, n = 8; cKO: 31.7 ± 4.7, n = 9; control *vs* cKO: p < 0.05, unpaired t-test) and increase in latency to stage 3 (Fig 4E, control: 21.2 ± 3.5, n = 8; cKO: 51.1 ± 13.4, n = 9; control *vs* cKO: p < 0.05, Mann Whitney test), respectively. Remarkably, c-Fos expression was significantly reduced in *Bdnf* cKO mice as compared to controls (Fig S2). These results indicate that seizure-induced MC-CG LTP may have a pro-convulsant action.

**Figure 4:**
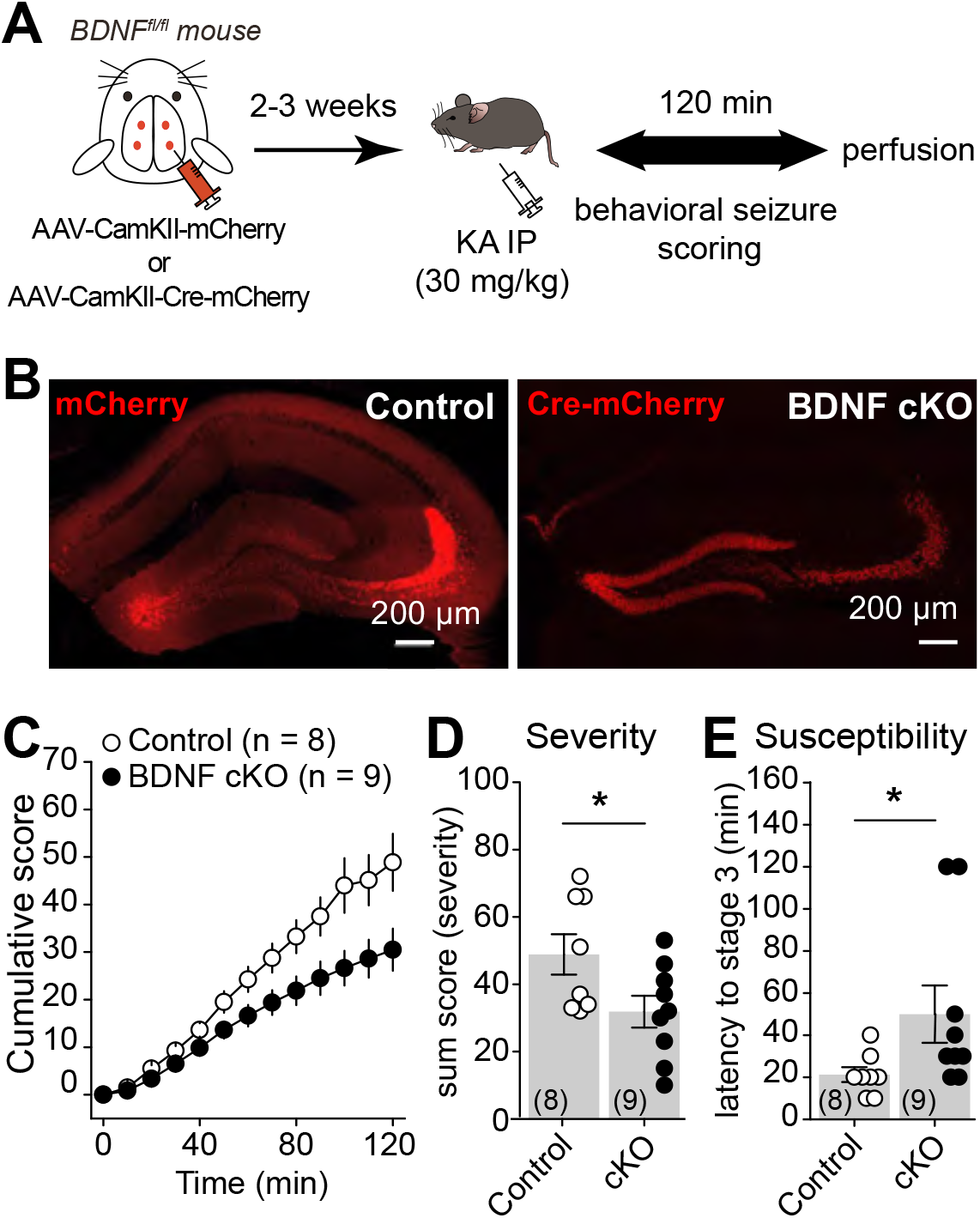
Knocking out BDNF from hippocampal excitatory neurons reduced KA-induced seizures. **(A)** AAV-CaMKII-mCherry (control) or AAV-CaMKII-Cre-mCherry (KO) was injected bilaterally into ventral and dorsal DG of *Bdnf*^fl/fl^ mice. **(B)** Confocal images (right) showing the viral expression in the DG. **(C-E)** Deletion of BDNF from hippocampal excitatory neurons (*Bdnf*^fl/fl^ mice injected with AAV-CaMKII-Cre-mCherry) induced a significant decrease in cumulative seizure score (C), in sum score (D) and a significantly increase in latency to convulsive seizures (E) as compared with controls (*Bdnf*^fl/fl^ mice injected with AAV-CaMKII-mCherry). * p < 0.05. Numbers in parentheses represent number of mice. Data are presented as mean ± SEM.

### *In vivo* application of MC-GC LTP induction protocol promoted epileptic seizures

We previously showed that optogenetic repetitive stimulation of MC axons triggers robust MC-GC LTP *in vitro*^24^. We therefore tested whether *in vivo* induction of LTP using optogenetic activation of MCs promotes seizure activity. To this end, we selectively expressed ChiEF in MCs by bilaterally injecting the Cre-recombinase-dependent AAV-hSyn-DIO-ChIEF-Tdtomato into the dorsal and ventral DG of *Drd2*-Cre mice (Fig 5A). Blue light stimulation (MC BS: 5 pulses of 5 ms at 30 Hz, repeated 50 times every 0.5 s) was delivered through an optic fiber placed above the dorsal DG IML *in vivo* and sham-light was used as a control (Fig 5A, B and F). Optic fiber location and selective viral expression in MCs were confirmed *posthoc* (Fig 5B). We also verified that the light stimulation protocol induced MC-GC LTP *in vivo*. If so, this plasticity should be occluded in hippocampal slices prepared after *in vivo* light stimulation. We found that repetitive light stimulation of MC axons failed to induce LTP *in vitro* when LTP protocol was pre-applied *in vivo* as compared to control slices (Fig 5C, after *in vivo* LTP: 96.1 ± 7.9 % of baseline, n = 6, p > 0.05, paired t-test; control: 146.7 ± 4.9 % of baseline, n = 5, p < 0.001, paired t-test; *in vivo* LTP *vs* control: p < 0.001, unpaired t-test). In addition, both PPR (Fig 5D, control: 1.51 ± 0.10, n = 5; *in vivo* LTP: 1.08 ± 0.09, n = 6; control *vs in vivo* LTP: p < 0.05, unpaired t-test) and CV (Fig 5E, control: 0.41 ± 0.04, n = 5; *in vivo* LTP: 0.25 ± 0.04, n = 6; control *vs in vivo* LTP: p < 0.05, unpaired t-test) were significantly reduced after *in vivo* photo-stimulation as compared to control mice (sham light). These results strongly suggest that *in vivo* light stimulation of MC axons (MC BS: 5 pulses of 5 ms at 30 Hz, repeated 50 times every 0.5 s) induced presynaptic LTP at MC-GC synapses. We then tested whether *in vivo* induction of MC-GC LTP can promote epileptic seizures. Seizures were induced with 20 mg/kg KA IP, 50 min after *in vivo* induction of LTP (Fig 5F). Light delivery alone was not sufficient to trigger any behavioral seizure (data not shown). Remarkably, we found that *in vivo* application of a MC-GC LTP induction protocol significantly increased the severity and susceptibility of convulsive seizures induced by KA, as indicated, respectively, by significant increase in cumulative seizure score (Fig 5G, two way ANOVA RM, light: F(1, 3) = 100.7, p < 0.001, time: F(1.3, 4.1) = 221.5, p < 0.001; light x time: F(1,3) = 109.3, p < 0.01), in sum score (Fig 5H, control: 18.0 ± 1.8, n = 5; *in vivo* LTP: 28.0 ± 1.5, n = 4; control *vs in vivo* LTP: p < 0.01, unpaired t-test), and a significantly decrease in latency to convulsive seizures (Fig 5I, control: 104.0 ± 10.3, n = 5; *in vivo* LTP: 42.5 ± 7.5, n = 4; control *vs in vivo* LTP: p < 0.05, Mann Whitney test). While optogenetic activation of MCs could impact their basic membrane properties or synaptic inputs, we found that the stimulation protocol that triggers LTP *in vivo* (5 MCs action potential at 30 Hz, repeated 50 times every 0.5 s) did not significantly alter MC membrane resistance nor the main excitatory synaptic drive, i.e., GC inputs, onto MC recorded *in vitro* (Fig S3) (see Methods). Taken together, these findings strongly suggest that *in vivo* induction of MC-GC LTP can worsen epileptic seizures.

**Figure 5:**
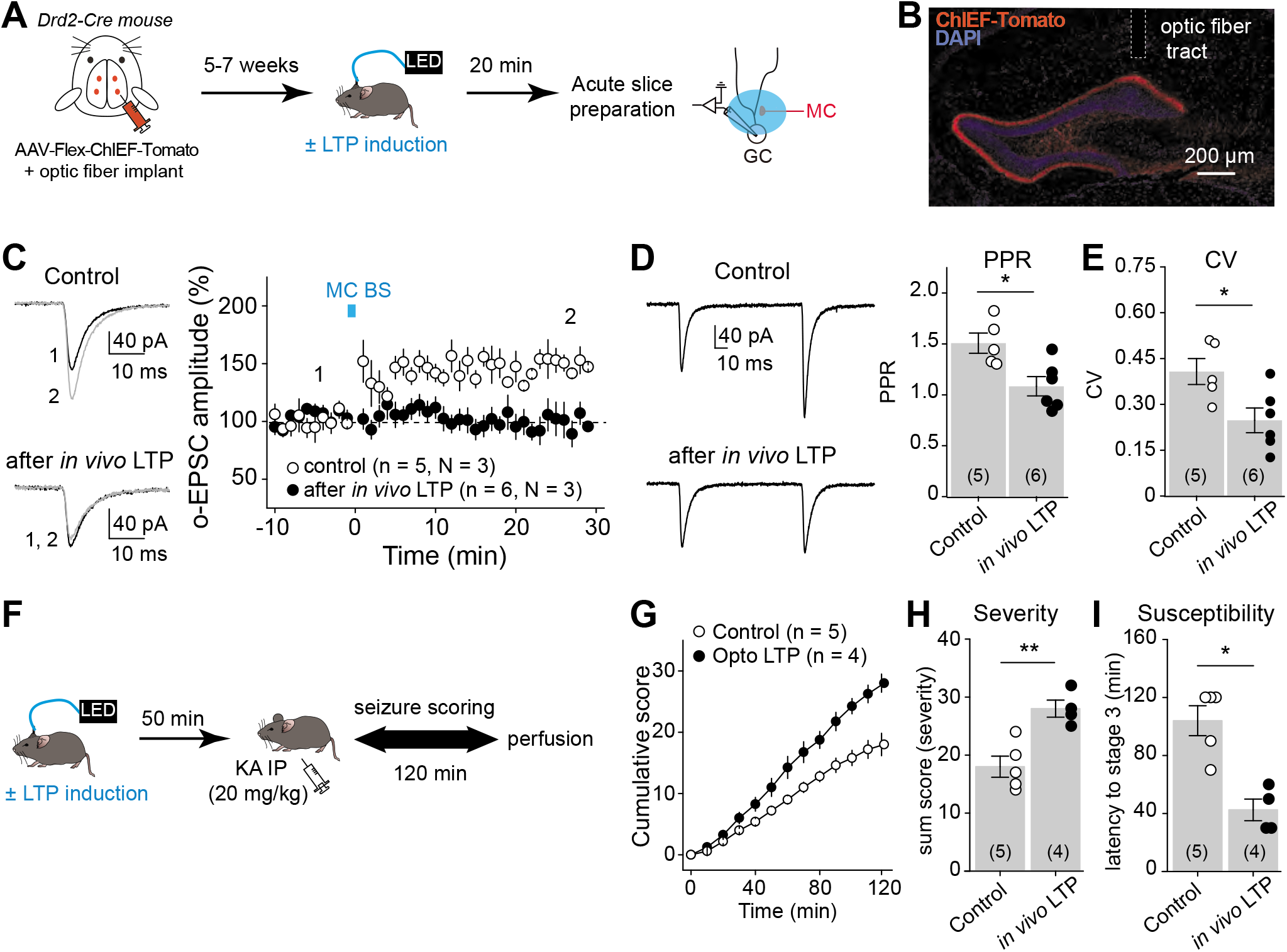
*In vivo* induction of MC-GC LTP promoted seizures. **(A)** Diagram of the experimental timeline. To selectively photo-stimulate MC axons *in vivo*, AAV-hSyn-ChiEF-Tdtomato was injected bilaterally into ventral and dorsal DG of Drd2-cre mice and an optical fiber was implanted above the IML of the DG. LTP induction protocol (MC BS: 5 pulses at 30 Hz, repeated 50 times, every 0.5 s) was applied *in vivo* by delivering blue light through a patch cord cable connected to a fiber-coupled 470 nm LED light source, 5-7 weeks after surgery. Sham-light was used as a control. Acute hippocampal slices were prepared 20 min later and MC-GC synaptic properties were analyzed using whole cell recordings of GCs and light stimulation of MCs axons. **(B)** Confocal images showing the viral expression and the fiber tract location. **(C)** O-EPSCs were recorded in GC in response to MC axon photostimulation. Representative traces (light) and time-course summary plot (light) showing that *in vivo* application of MC BS prevented subsequent induction of LTP *in vitro*. Numbers in parentheses, indicate the number of cells (n) and the number of mice (N). **(D, E)** PPR (D) and CV (E) were both significantly decreased after *in vivo* application of MC-GC LTP induction protocol, as compared to sham light (control) condition. Numbers in parentheses, indicate the number of cells. **(F)** Diagram of the experimental timeline. LTP induction protocol (MC BS: 5 pulses at 30 Hz, repeated 50 times, every 0.5 s) was delivered *in vivo* while sham-light was used as a control. KA (20 mg/kg IP) was then administered 50 min after the optogenetic stimulation and seizures were scored for 120 min. **(G-I)** Application of MC-GC LTP induction protocol increased seizure severity and susceptibility as indicated, respectively, by significant increase in cumulative seizure score (G), in sum score (H), and a significantly decrease in latency to convulsive seizures (I). * p < 0.05, ** p < 0.01. Data are presented as mean ± SEM.

## DISCUSSION

In this study, we found that early seizures potentiate crucial hippocampal excitatory synapses, thereby facilitating further epileptic activity. KA-induced acute epileptic seizures, not only increased MC and GC activity *in vivo*, but also triggered a BDNF-dependent strengthening of MC-GC synaptic transmission that occluded subsequent induction of MC-GC LTP. In addition, blocking MC-GC LTP and silencing MC selectively, were both associated with significant decrease in seizure susceptibility and severity. Moreover, *in vivo* induction of MC-GC LTP was sufficient to worsen convulsive seizures subsequently triggered with KA. Overall, our findings strongly suggest that seizure-induced plasticity at MC-GC excitatory synapses may promote epilepsy and significantly contribute to the pro-convulsant role of MCs during early stages of epilepsy.

### Initial seizures induced BDNF-dependent strengthening of MC-GC synaptic transmission

Using *in vivo* 2-photon live imaging in awake behaving mice, we found that acutely induced seizures triggered a massive increase in both MC and GC calcium signal (Fig 2), indicating a robust increase in neuronal activity. Our findings are consistent with a recent study reporting *in vivo* epileptiform calcium signals detected with fiber photometry in the DG following KA administration^35^. While these signals likely reflect the activity of a large population of neurons, including interneurons, we could assess calcium activity of individual MCs and GCs by combining selective expression of a calcium indicator in DG excitatory neurons and 2-photon live imaging. It has been reported that dorsal GCs are mainly activated by ventral MCs^34^. However, because of the limited access to the ventral hippocampus, 2-photon live imaging did not allow us to directly assess which of MCs or GCs take the lead during epileptic activity. We hypothesize that KA administration activates MCs which in turn engage GCs. Consistent with this idea, we have recently found that MCs express functional extrasynaptic kainate receptors whose activation with sub-micromolar concentrations of KA can drive MC activity *in vitro*; whereas GCs show comparatively much less sensitivity (at least one order of magnitude) to KA application^38^. Although interneurons^39^ and CA3 pyramidal cells also express kainate receptors^40,41^, it is unlikely that activation of these neurons could directly drive GCs. Furthermore, MCs show higher activity *in vivo* in contrast to GCs^42–44^, making them more likely to be engaged during epileptic activity, independently of the nature of the chemoconvulsant. This last notion is also supported by the fact that MC silencing not only reduced KA-induced seizures (Fig 1) but also prevents pilocarpine-induced epilepsy^5^. Although MCs also excite inhibitory interneurons^17^, the anti-convulsant effect of MC silencing (Fig 1) suggests that MC silencing during initial seizures has a stronger impact on the activity of GCs than interneurons^5^. Besides inducing synaptic plasticity, KA-induced increase in MC and GC intracellular calcium concentration (Fig 2) may also contribute to excitotoxicity and cell death. Altogether, our findings demonstrate that MCs and GCs are highly active during initial experimental seizures, suggesting that sustained activation of MCs contributed to GC recruitment.

We gathered multiple lines of evidence indicating that initial convulsive seizures induced presynaptically expressed MC-GC LTP *in vivo*. MC-GC synaptic strength was increased in KA-treated mice as compared to sham-injected animals, and this strengthening was associated with a significant reduction in both PPR and CV (Fig 2), suggesting a presynaptic mechanism. Furthermore, induction of MC-GC LTP *in vitro* was occluded after convulsive seizures (Fig 2), indicating a common step. The KA-induced strengthening of MC-GC synaptic transmission *in vivo* was likely induced by the increase in MC activity^38^, consistent with the observation that repetitive MC activity triggers robust MC-GC LTP in acute rodent hippocampal slices^24^. Although *in vitro* epileptic activity was associated with a rise in the net excitatory drive between MCs and GCs^5^, it is unclear whether this effect results from disinhibition or direct MC-GC synaptic strengthening. Our findings show that both *in vivo* optogenetic activation of MCs (Fig 5) and acute seizures (Fig 2) were sufficient to trigger presynaptic MC-GC LTP.

Several studies indicate that seizures can increase both BDNF levels^45–48^ and TrkB activation in the hippocampus^26,49^. In addition, BDNF is necessary and sufficient for MC-GC LTP^24^, and it can be released from both MCs and GCs following MC repetitive activity *in vitro*^25^. It is therefore likely that by releasing BDNF, MC and GC activity induces MC-GC LTP *in vivo*. In support of this mechanism, we found that genetic removal of *Bdnf* from GCs abolished seizure-induced MC-GC LTP (Fig 3), while it did not affect basal MC-GC synaptic (Fig 3) nor GC membrane properties in sham-injected mice (Fig S1). Of note, we did not observe any failure of seizure induction when *Bdnf* was sparsely KO. Altogether, our new findings indicate that BDNF mediates *in vivo* seizure-induced strengthening of MC-GC excitatory synapses.

### Seizure-induced LTP at MC-GC synapse promotes epilepsy

Our findings strongly suggest that activity-dependent strengthening of MC-GC synapses promotes acute seizures. While MCs innervate GCs and inhibitory interneurons, MC repetitive activity that induces MC-GC LTP, at least *in vitro*, has no effect on feed-forward inhibition onto GCs^24^, and such repetitive activity does not induce plasticity at GC-MC synapses either (Fig S3). LTP-induced worsening of seizures (Fig 5) is supported by the extensive MC projection onto the proximal dendrites of GCs^17^ and the powerful MC-GC excitatory drive reported *in vitro*^24^ and *in vivo*^34^. Given that a single MC innervates as much as 75% of the septotemporal axis of the hippocampus^50^, broad induction of MC-GC LTP can be detrimental, underscoring a link between uncontrolled LTP at hippocampal excitatory synapse and seizures. Conversely, blocking activity-dependent strengthening of MC-GC synapses *in vivo* reduced seizures. Protein kinase A (PKA) and BDNF signaling pathways are both necessary and sufficient for activity-dependent LTP at MC-GC synapses^24,25^. MC silencing using G_i_ inhibitory DREADD (Fig 1), and knocking out *Bdnf* from hippocampal excitatory neurons (Fig 4) both reduced acute KA-induced seizures. However, MC silencing not only prevents MC-GC LTP induction but also reduces basal MC-GC synaptic transmission (Fig 1) and MC output activity. Although *Bdnf* cKO had no effect on MC-GC basal transmission (Fig 3) and GC excitability (Fig S1), we cannot discard an effect on other synapses including GC-CA3 synaptic transmission and plasticity. *Bdnf* deletion reduced c-Fos expression in the DG of KA-injected mice (Fig S2), suggesting that BDNF/TrkB signaling contributes to seizure-induced opening of the DG gate likely by strengthening MC-GC synapses. Consistent with this scenario, optogenetic induction of MC-GC LTP *in vivo* was sufficient to worsen convulsive seizures. Notably, type-1 cannabinoid receptors, which are highly expressed at MC terminals^51^, tonically suppress MC-GC transmission and also dampen the induction of MC-GC LTP^52^ in an activity-dependent manner. By suppressing excitatory drive, these receptors could be a potential target to prevent epilepsy^53–55^. In agreement with recent findings using the pilocarpine-model^5^, our results strongly support a pro-convulsant role of MCs during early epilepsy. In contrast, in a chronic mouse model of TLE induced by KA intrahippocampal administration, MCs are reportedly anti-epileptic^4^, suggesting that the role of MCs may differ significantly with the disease stage.

Compelling evidence indicates that BDNF and its high-affinity receptor TrkB promote worsening of TLE^26,27,36,37,56–60^, but the precise mechanisms and specific contribution of different cell types are not entirely clear. Here we identified two reciprocally connected excitatory neurons in the dentate gyrus, MCs and GCs, that can mediate the BDNF pro-convulsant effects via BDNF-dependent MC-GC LTP. Interfering with BDNF/TrkB signaling in different ways reduced epilepsy--i.e. heterozygous deletion of *Bdnf*^36^, neuronal deletion of *Bdnf* or *TrkB*^37^, chemogenetic blockade of TrkB kinase activity in TrkB^F616A^ mutant mice^59^ and a mutant mouse that uncouples TrkB from its downstream phospholipase Cγ1 signaling^57^. Importantly, based on our previous^24^ and present findings (Fig 3,5), these manipulations could also prevent seizure-induced MC-GC LTP and the associated facilitation of epileptic activity. Conversely, overexpression of BDNF in the brain^60^ and local infusion of BDNF into the hippocampus^56^ worsen epileptic seizures. Because BDNF is sufficient to induce MC-GC LTP *in vitro*^24^, it is likely that *in vivo* infusion of BDNF promotes seizures by inducing MC-GC LTP broadly. Of note, exogenous BDNF delivery into the hippocampus of chronically epileptic rats can have anti-epileptic and neuroprotective effects^61^, suggesting that BDNF action might differ with epilepsy stages.

Altogether our findings uncover a potential mechanism implicated in the early stages of epilepsy, before the brain damages become irreversible. We highlighted how initial seizures can shape an important but overlooked hippocampal excitatory synapses in a BDNF-dependent manner, and how broad, uncontrolled induction of LTP can be detrimental and promote subsequent induction of seizures. Manipulations that suppress LTP induction and BDNF signaling at MC-GC synapses may be a new strategy for the treatment of epilepsy.

## METHODS

### Experimental Model and Subject Details

C57BL/6, *Bdnf* floxed (*Bdnf*^fl/fl^ or *Drd2*-cre (B6.FVB(Cg)-Tg(Drd2-cre)ER44Gsat/Mmucd, MMRRC 032108-UCD) mice (2-3.5–month-old, both males and females) were used in this study. All animals were group housed in a standard 12 hr light/12 hr dark cycle and had free access to food and water. Animal handling, breeding and use followed a protocol approved by the Animal Care and Use Committee of Albert Einstein College of Medicine, in accordance with the National Institutes of Health guidelines. *Bdnf*^fl/fl^ mice, generated by Dr. Jaenisch, were kindly donated by Dr. Lisa Monteggia (University of Texas, Southwestern Medical Center).

### Acute hippocampal slice preparation

Acute transverse hippocampal slices (300 μm thick) were prepared from C57BL/6, *Bdnf*^fl/fl^ and *Drd2*-cre mice. Animals were anesthetized with isoflurane and euthanized in accordance with institutional regulations. The hippocampi were then removed and cut using a VT1200s Microslicer (Leica Microsystems Co.) in a solution containing (in mM): 93 N-methyl-D-glucamine (NMDG), 2.5 KCl, 1.25 NaH_2_PO_4_, 30 NaHCO_3_, 20 HEPES, 25 D-glucose, 2 Thiourea, 5 Na-Ascorbate, 3 Na-Pyruvate, 0.5 CaCl_2_, 10 MgCl_2_, 93 HCl. These slices were then transferred to 32°C extracellular artificial cerebrospinal fluid (ACSF) solution, containing (in mM): 124 NaCl, 2.5 KCl, 26 NaHCO_3_, 1 NaH_2_PO_4_, 2.5 CaCl_2_, 1.3 MgSO_4_ and 10 D-glucose, for 30 min and then kept at room temperature for at least 40 min before recording. All solutions were equilibrated with 95% O_2_ and 5% CO_2_ (pH 7.4).

### Electrophysiology

All recordings were performed at 28 ± 1°C in a submersion-type recording chamber perfused at 2 ml/min with ACSF. GABA_A_ and GABA_B_ receptor antagonists, picrotoxin (100 μM) and CGP55845 hydrochloride (3 μM), were included in the extracellular solution (ACSF) except in the experiments shown in Fig S3. Whole-cell patch-clamp recordings using a Multiclamp 700A amplifier (Molecular Devices) were made from GCs (in the upper blade of the DG) and MCs (Fig S3) voltage clamped at −60 mV (V_h_ = −60 mV) using patch-type pipette electrodes (~3-4 MΩ) containing (in mM): 135 KMeSO_4_, 5 KCl, 1 CaCl_2_, 5 NaOH, 10 HEPES, 5 MgATP, 0.4 Na_3_GTP, 5 EGTA and 10 D-glucose, pH 7.2 (288-291 mOsm). Series resistance (~6-25 MΩ for GCs and ~17-21 MΩ for MCs) was monitored throughout all experiments with a −5 mV, 80 ms voltage step, and cells that exhibited a significant change in series resistance (> 20%) were excluded from analysis. Mature GCs were identified by characteristic hyperpolarized membrane potential (checked immediately after membrane break in, −72 to −83 mV). Cells with high membrane resistance (> 800 MΩ) were considered as putative adult born GCs^62^ and were excluded from the analysis. To activate MC axons, a broken tip (~10–20 μm) stimulating patch-type micropipette filled with ACSF was placed in the IML (< 50 μm from the border of the GC body layer) and paired, monopolar square-wave voltage or current pulses (100 μs pulse width, 4-30 V) were delivered through a stimulus isolator (Digitimer DS2A-MKII). Typically, stimulation intensity was adjusted to obtain comparable magnitude of synaptic responses across experiments, e.g., 30-80 pA EPSCs (V_h_ = −60 mV), except for input/output experiments shown in Fig 2F, 3B and 3D. MC-GC LTP was typically induced by 5 stimuli at 100 Hz repeated 50 times, every 0.5 s, except in Fig 5 (see below). Light evoked EPSCs (o-EPSCs), shown in Fig 5 C-E, were triggered using 1 ms pulses of blue light, provided by a collimated LED (Thorlabs, M470L3-C5, 470 nm, 300mW) and delivered through the microscope objective (40X, 0.8 NA). Recordings were performed in acute hippocampal slices < 700 μm from the optic fiber implant. The light was centered in the IML, and the light intensity adjusted to obtain ~60–120 pA responses (V_h_ = −60 mV). In the experiments shown in Fig 5C, LTP was induced using light stimulation of MC axons with the following protocol: 5 pulses of 1 ms, at 30 Hz repeated 50 times, every 0.5 s. In Fig S3, GC-MC synaptic strength was monitored. MCs identity was confirmed by the presence of a characteristic high frequency of spontaneous EPSCs, and by checking the firing pattern in response to depolarizing step of currents (non-burst firing and action potentials with almost no afterhyperpolarization)^63^. GC axons were electrically stimulated in the DG subgranular zone using a bipolar theta glass ACSF-filled pipette. The mGluR2/3 agonist DCG-IV (1 μM), which selectively reduces GC-MC synaptic transmission^64,65^ was applied at the end of each experiment to confirm the nature of the stimulated input. To mimic physiological conditions, EPSCs were recorded in absence of GABA receptor blockers. Reagents were bath applied following dilution into ACSF from stock solutions stored at −20°C prepared in water or DMSO, depending on the manufacturer’s recommendation.

### BDNF conditional KO

*Bdnf*^fl/fl^ mice were injected with Cre expressing (AAV5-CamKII-Cre-mCherry, 5.8 x10^12^ Virus Molecules/mL, UNC Vector) or control (AAV5-CamKII-mCherry, 4.9 x10^12^ Virus Molecules/mL, UNC Vector) adeno-associated viruses. For electrophysiological experiments, 0.5 μl of virus was injected (flow rate of 0.1 μl/min) unilaterally into the dorsal blade of the DG (relative to bregma: 2.06 mm posterior, 1.5 mm lateral, 1.8 mm ventral) of 5-8–week-old (w.o.) *Bdnf*^fl/fl^ mice. Slices for electrophysiology were prepared from injected animals, 2 to 4 weeks after injection. In postsynaptic cKO animals, we verified the absence of mCherry in the hilus of the whole ipsilateral hippocampus, as previously described^24^. For seizure monitoring and c-Fos labeling experiments shown in Fig 4 and Fig S2, 0.5 μl of virus was injected (flow rate of 0.1 μl/min) bilaterally into both the dorsal (relative to bregma: 1.9 mm posterior, 1.25 mm lateral, 2.1 mm ventral) and ventral (relative to bregma: 3.2 mm posterior, 2.2 mm lateral, 2.8 mm ventral) DG of 5-8–w.o. *Bdnf*^fl/fl^ mice and seizure was induced and monitored 2-3 weeks post-injection. Animals were placed in a stereotaxic frame and anesthetized with isoflurane (up to 5% for induction and 1%–3% for maintenance). In all experiments, both male and female mice were used with a similar ratio for the two types of viruses.

### MC silencing

To silence MCs, the Gi inhibitory DREADD (iDREADD) was selectively expressed in MCs. The Cre-dependent AAV-CamKII-DIO-hM4D(G_i_)-mCherry (4.16 x 10^13^ vg/mL, prepared at Janelia Research Campus) or AAV-CamKII-DIO-mCherry (control, 5.95 x10^13^ vg/mL, prepared at Janelia Research Campus) was injected (0.5 μl/site, at 0.1 μl/min) bilaterally into both the dorsal (relative to bregma: 1.9 mm posterior, 1.25 mm lateral, 2.1 mm ventral) and ventral (relative to bregma: 3.2 mm posterior, 2.2 mm lateral, 2.8 mm ventral) DG of 5-8–w.o. *Drd2*-Cre mice. Seizures were induced and monitored 2-3 weeks post-injection. The DREADD selective agonist clozapine-N-oxide (CNO, TOCRIS) was bath applied in acute slices in experiments shown in Fig 1D. CNO was administered intraperitoneally in Fig 1E-H (2 mg/kg diluted in saline solution containing 2% DMSO) 30 min before *in vivo* seizure induction (KA IP), based on kinetics of CNO plasma levels and *in vivo* effects ^66,67^. Viral expression was verified *posthoc*.

### *In vivo* induction of MC-GC LTP with optogenetics

To induce MC-GC LTP *in vivo* by light-activating MCs, ChiEF was selectively expressed in MCs, and an optic fiber (200 μm diameter) was implanted unilaterally (relative to bregma: 1.9 mm posterior, 1.25 mm lateral, 1.8 mm ventral) to deliver blue light above the IML of the DG. The Cre-dependent AAV-hSyn-Flex-ChIEF-Tdtomato (2.4 x10^12^ Virus Molecules/mL) was injected bilaterally (0.5 μl/site, at 0.1 μl/min) into both the dorsal (relative to bregma: 1.9 mm posterior, 1.25 mm lateral, 2.1 mm ventral) and ventral (relative to bregma: 3.2 mm posterior, 2.2 mm lateral, 2.8 mm ventral) DG of 5-8–w.o. *Drd2*-Cre mice. 5-7 weeks after surgery, MC-GC LTP induction protocol^24^ was applied *in vivo*, by delivering a brief burst of blue light (5 pulses of 5 ms at 30 Hz, repeated 50 times every 0.5 s), using a fiber-coupled 470 nm LED light source (Thorlabs, MF470F3, ~7 mW output from the 200 μm optic fiber). Viral expression and optic fiber implant location were confirmed *posthoc*.

### Seizure induction and monitoring

Seizures were induced acutely using intraperitoneal (IP) injection of 20-30 mg/kg of kainic acid (KA, HelloBio HB0355), prepared in saline solution the same day, in 2-3–month-old mice. To be able to see potential decrease and increase in seizure severity/susceptibility, 30 mg/kg (Fig 1 and 4) and 20 mg/kg (Fig 5) of KA were used, respectively. Of note, seizures were less severe in *Drd2*-Cre mice as compared to C57BL/6 (wild type) and *Bdnf*^fl/fl^ animals, likely due to strain differences^33^. For MC-GC basal transmission and synaptic plasticity analysis, acute hippocampal slices were prepared when the animal reached stage 3 of convulsive seizures, i.e., forelimb clonus and rearing (Fig 2E-I) or 25 min post KA IP injections (Fig 3). Saline-injected mice were used as a control. For behavioral seizure scoring, mice were monitored during 120 min post-injection and behavioral seizures were scored, by an experimenter blind to condition (control *vs Bdnf* cKO or control *vs* MC silencing), using a modified Racine scale^68^ as follows: stage 0: normal behavior, stage 1: immobility and rigidity, stage 2: head bobbing, stage 3: forelimb clonus and rearing, stage 4: continuous rearing and falling, stage 5: clonic-tonic seizure, stage 6: death. The maximum Racine score was recorded every 10 minutes and the cumulative seizure score was obtained by summing these scores across all 12 bins of the 120 min experiment. Mouse movements were visualized with an infrared camera for 2P imaging experiments (see below). For MC silencing experiments, IP injection of CNO (2 mg/kg, I.P) was delivered 30 min before inducing seizures with 30 mg/kg KA IP injection to both groups, control and inhibitory DREADD. For in vivo optogenetic experiments, LTP was induced 50 min before KA IP injection, and animals receiving sham-light stimulation were used as control.

### c-Fos immunolabeling and posthoc confirmation of AAV expression

Two hours after KA (20-30 mg/kg in saline) intraperitoneal injection, mice were deeply anesthetized using isoflurane (3-5%) and transcardially perfused with 4% paraformaldehyde (PFA) in 0.1 M sodium phosphate buffer (PBS). After 24-48h fixation in 4% PFA, 50 μm-thick brain coronal sections were prepared using a DSK Microslicer (DTK-1000). Brain slices were washed in PBS (3 x 10 min) and then incubated for 2h at room temperature in a blocking solution containing 10% goat serum, 1% Triton X-100 in PBS. Primary antibody was applied for 24 h, at 4°C (rabbit anti-c-Fos, 1:1000; Cell Signaling, cat # 2250S, 0.1% Triton X-100, 5% goat in PBS). After 4 washes of 10 min in PBS, the sections were incubated with AlexaFluor 488-conjugated goat anti-rabbit (1:500, Invitrogen, cat # A11008) for 2 h at room temperature. Slices were washed twice in PBS for 10 min each, stained with 40,6-diamidino-2-phenylindole (DAPI, 1:1000 in PBS, 20 min, ThermoFisher) to label cell nuclei and mounted with Prolong diamond antifade reagent mountant (ThermoFisher) onto microscope slides.

### Image Acquisition and cell quantification

Images were acquired using a Zeiss LSM 880 Airyscan Confocal microscope with Super-Resolution and ZEN (black edition) software and a 25X oil-immersion objective. All images were analyzed using Fiji and cells were counted using Cell Counter plug-in in Fiji, in the upper and lower blade of the DG in areas of maximum viral expression (> 70% of mCherry+ neurons), by an experimenter blind to the conditions. All infected cells were confirmed by using the DAPI channel, and individual counts were taken for c-Fos-expressing neuron to provide a percentage of c-Fos+ cells among the total number of neurons. For each animal, we used 1 dorsal and 1 ventral section. Right and left hippocampus were randomly used.

### Adeno-Associated Virus Vector Construction

For the construction of AAV plasmids, the human Synapsin (hSyn) promoter in pAAV-hSyn-DIO-hM4D(Gi)-mCherry (Addgene, cat #: 50459) was replaced with mouse CaMKIIa promoter using MluI and SalI restriction sites to produce pAAV-CaMKII-DIO-hM4D(Gi)-mCherry and pAAV-CaMKII-DIO-mCherry. The AAVs were prepared at Janelia Research Campus.

### Data analysis

Electrophysiological data were acquired at 5 kHz, filtered at 2.4 kHz, and analyzed using a Multiclamp 700A amplifier (Molecular Devices) and custom-made software for IgorPro 7.01 (Wavemetrics Inc.). PPR was defined as the ratio of the amplitude of the second EPSC (baseline taken 1-2 ms before the stimulus artifact) to the amplitude of the first EPSC. The interval between first and second EPSC was 100 ms. CV was calculated as the standard deviation of EPSC amplitude divided by mean EPSC amplitude. For each experiment, an average of 30 consecutive PPRs and CVs were used. The magnitude of LTP was determined by comparing 10 min baseline responses with responses 20-30 min after induction protocol. Averaged traces include 20 consecutive individual responses (or 5 consecutive responses for input/output experiments in Figures 2F, 3B and 3D).

### In vivo 2-photon imaging

WT mice (6-8–w.o.) were injected in the right dorsal hippocampus (relative to bregma: 1.2 mm posterior, 1.6 mm lateral, 1.9 mm ventral) with a viral vector encoding for the red-shifted Ca^2+^ sensor jRGECO1a [AAVDJ-CaMKIIa::NES-jRGECO1a-WPRE-SV40 (AAV-CaMKII-jRGECO1a), 950μL at 5.4 x10^12^ viral particles/mL, University of North Carolina Vector Core, plasmid kindly donated by Dr. Fred Gage]. A 3-mm diameter craniotomy was drilled around the injection site 24 hrs after viral injection^69^, the meninges, cortex, and corpus callosum were removed by aspiration and the hippocampus and alveus fibers were left intact. The lesion was irrigated with sterile saline throughout the procedure. A titanium window (3-mm diameter, 1.3-mm deep) with a glass bottom was implanted above the hippocampus. The implant was secured using dental cement and a titanium bar (34 × 4 × 1.3 mm) was attached to head-fix the mice to the microscope set-up. *In vivo* calcium imaging was performed 3-4 weeks after surgery, using a two-photon microscope (Thorlabs Bergamo) equipped with a 16x 0.8NA objective (Nikon) and a Fidelity-2 1070 nm laser (Coherent). The mice were head-fixed and placed on a 180 cm-long treadmill belt. For optimum light transmission, the angle of the mouse’s head was adjusted to ensure that the imaging window was parallel to the objective. Movies of Ca^2+^ activity were acquired at 15 frames/s using an average laser power of ~180 mW, as measured in front of the objective.

Stage 3 convulsive seizures (forelimb clonus) were identified using an infra-red camera. Raw calcium movie data was preprocessed using Suite2p^70^ applying default settings for alignment, registration, and extraction of regions of interest (ROIs). Manual curation was then performed to select significant ROIs according to morphology and activity data. Raw calcium traces and neuropil were extracted for further analysis with a custom written algorithm (Python). The neuropil was subtracted from each detected cell to remove contamination. For KA-injected mice, the mean fluorescence was determined for 500 frames selected during the onset of behavioral seizures (as determined by the observation of forelimb clonus), for each cell. For baseline activity, 500 frames were selected before IP injection. For control groups, frame selection was performed similarly before (baseline) and after saline injection, matching the time of KA injected mice. ΔF/F was calculated as follows for both groups, for each cell:

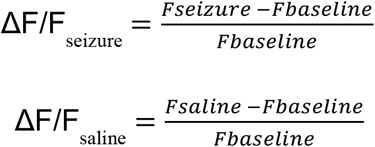

### Quantification and Statistical Analysis

The normality of all distributions was assessed using the Shapiro-Wilk test. In normal distributions, Student’s unpaired and paired two-tailed t tests were used to assess between-group and within-group differences, respectively. The non-parametric paired sample Wilcoxon signed rank test and Mann-Whitney’s U test were used in non-normal distributions. Statistical comparison in Fig 1F, 2F, 3B, 3D, 4C, 5G was performed using two-way ANOVA with repeated measure (RM) and Greenhouse-Geiser was used for correction of degrees of freedom when sphericity was not assumed. Statistical significance was set to p < 0.05 (***indicates p < 0.001, ** indicates p < 0.01, and * indicates p < 0.05). All values are reported as the mean ± SEM. Experiments shown in Fig 1E-H, Fig 3, Fig S1-2 and Fig 4 were performed in blind manner during data acquisition and analysis. Statistical analysis was performed using OriginPro (version b9.2.272) software (OriginLab).

## ACKNOWLEDGMENTS

We thank the members of the Castillo lab for constructive feedback. We also thank Pascal Kaeser (Harvard University) for sharing an AAV-hSyn-Flex-ChIEF-TdTomato plasmid and Lisa Monteggia (Vanderbilt University) for sharing *Bdnf*^fl/fl^ mice. This research was supported by the National Institutes of Health (NIH), R01-NS113600, R01-MH125772; R01 and MH125772 to P.E.C.; Fondation pour la Recherche Médicale (Postdoctoral Fellowship for a research abroad), the Fondation Bettencourt Schueller (Prix pour les Jeunes Chercheurs 2016) and the American Epilepsy Society Postdoctoral Research Fellowship (2020) to K.N; The Einstein Training Program in Stem Cell Research from the Empire State Stem Cell Fund through New York State Department of Health Contract C34874GG to M.A.F.; and a Whitehall Foundation Research Grant (2019-05-71) to J.T.G.

## Author contributions

K.N. and P.E.C. designed research and wrote the manuscript; K.N. performed and analyzed all experiments except the 2P imaging experiments that were performed and analyzed by M.A.F. and immunostainings that were performed and quantified by S.P. All authors edited the manuscript.

## The authors declare no competing interests

**Figure S1 (related to Figure 3):**
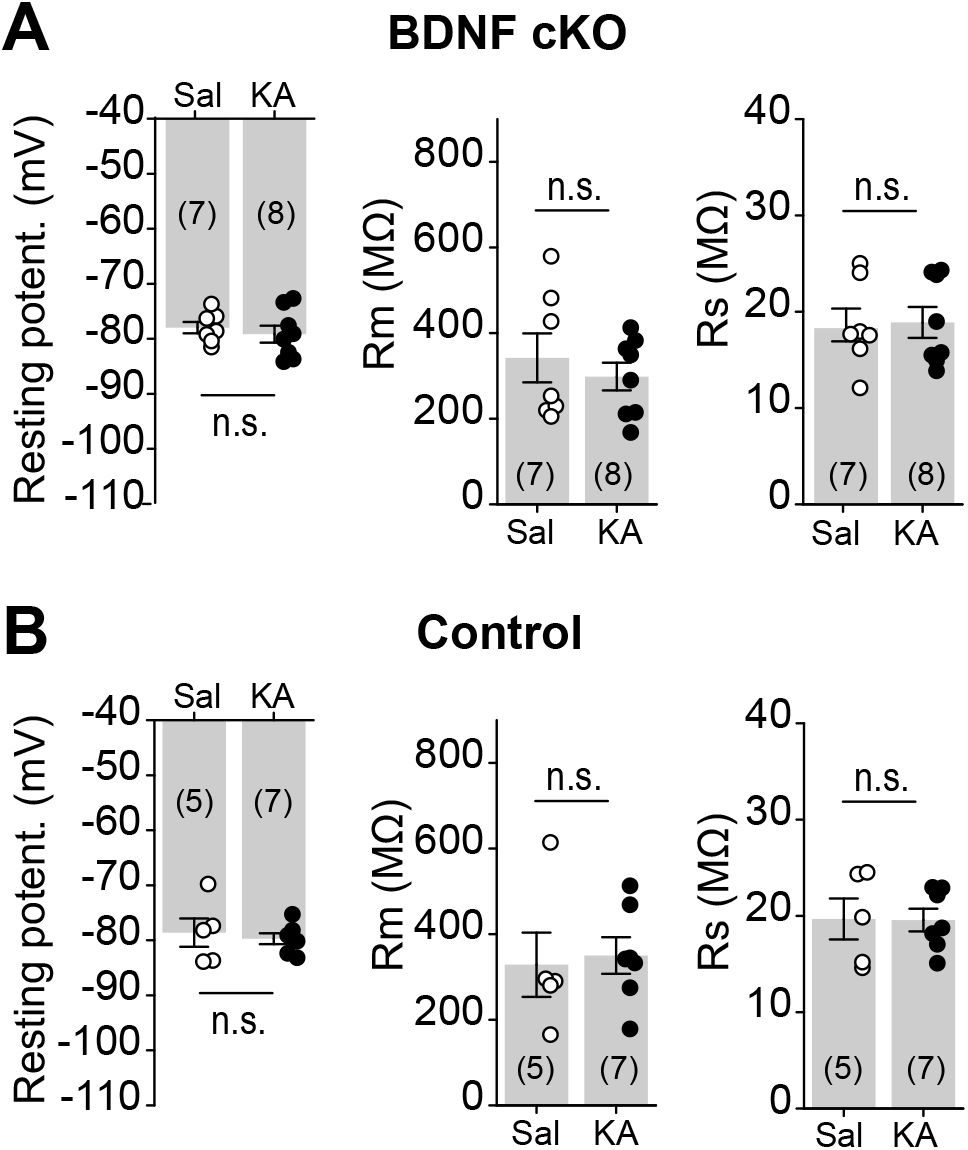
BDNF cKO did not alter GC membrane properties. **(A)** *Bdnf^fl/fl^* mice were injected with Cre-expressing virus (AAV-CaMKII-Cre-mCherry, BDNF cKO). Whole cell recordings were performed in Cre-mCherry-expressing GCs. Resting potential (saline: −77.3 ± 1.0 mV, n = 7; KA: −79.2 ± 1.6 mV, n = 8; saline vs KA: p > 0.05, unpaired t-test), membrane resistance (Rm, saline: 340.9 ± 57.0 MΩ, n = 7; KA: 298.5 ± 32.9 MΩ, n = 8; saline vs KA: p > 0.05, unpaired t-test) and series resistance (Rs, saline: 18.3 ± 1.7 MΩ, n = 7; KA: 18.9 ± 1.6 MΩ, n = 8; saline vs KA: p > 0.05, unpaired t-test) were not different in KA vs saline-treated. **(B)** *Bdnf*^fl/fl^ mice were injected with a control virus (AAV-CaMKII-mCherry). Membrane properties were similar in KA- and saline-injected mice (Resting potential, saline: −78.6 ± 2.6 mV, n = 5; KA: −79.7 ± 1.0 mV, n = 7; saline vs KA: p > 0.05, unpaired t-test; Rm, saline: 328.7 ± 75.2 MΩ, n = 5; KA: 350.2 ± 42.5 MΩ, n = 7; saline vs KA: p > 0.05, Mann Whitney test; Rs, saline: 19.7 ± 2.1 MΩ, n = 5; KA: 19.6 ± 1.8 MΩ, n = 7; saline vs KA: p > 0.05, unpaired t-test).

**Figure S2 (related to Figure 4):**
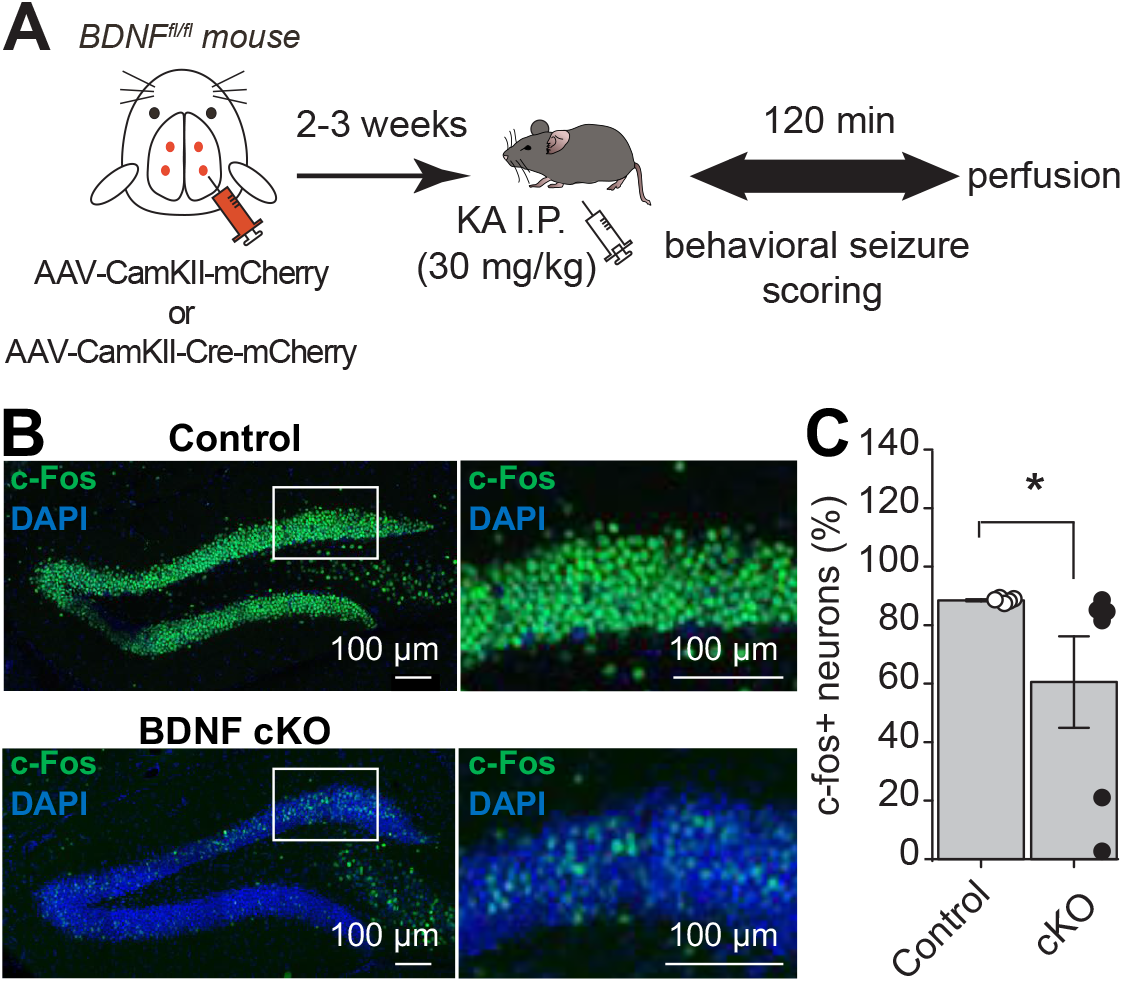
Neuronal BDNF cKO protected DG gating properties. **(A)** Diagram of the experimental timeline. **(B and C)** Confocal images (B) and quantification (C) showing how BDNF KO reduced the number in c-fos+ neurons in the DG of KA-injected mice (control: 88.5 ± 0.3% of c-Fos+ neurons, N =5 mice; BDNF cKO: 60.5 ± 15.6% of c-Fos+ neurons, N = 5 mice; control *vs* BDNF cKO: p < 0.05, Mann Whitney test).

**Figure S3 (related to Figure 5):**
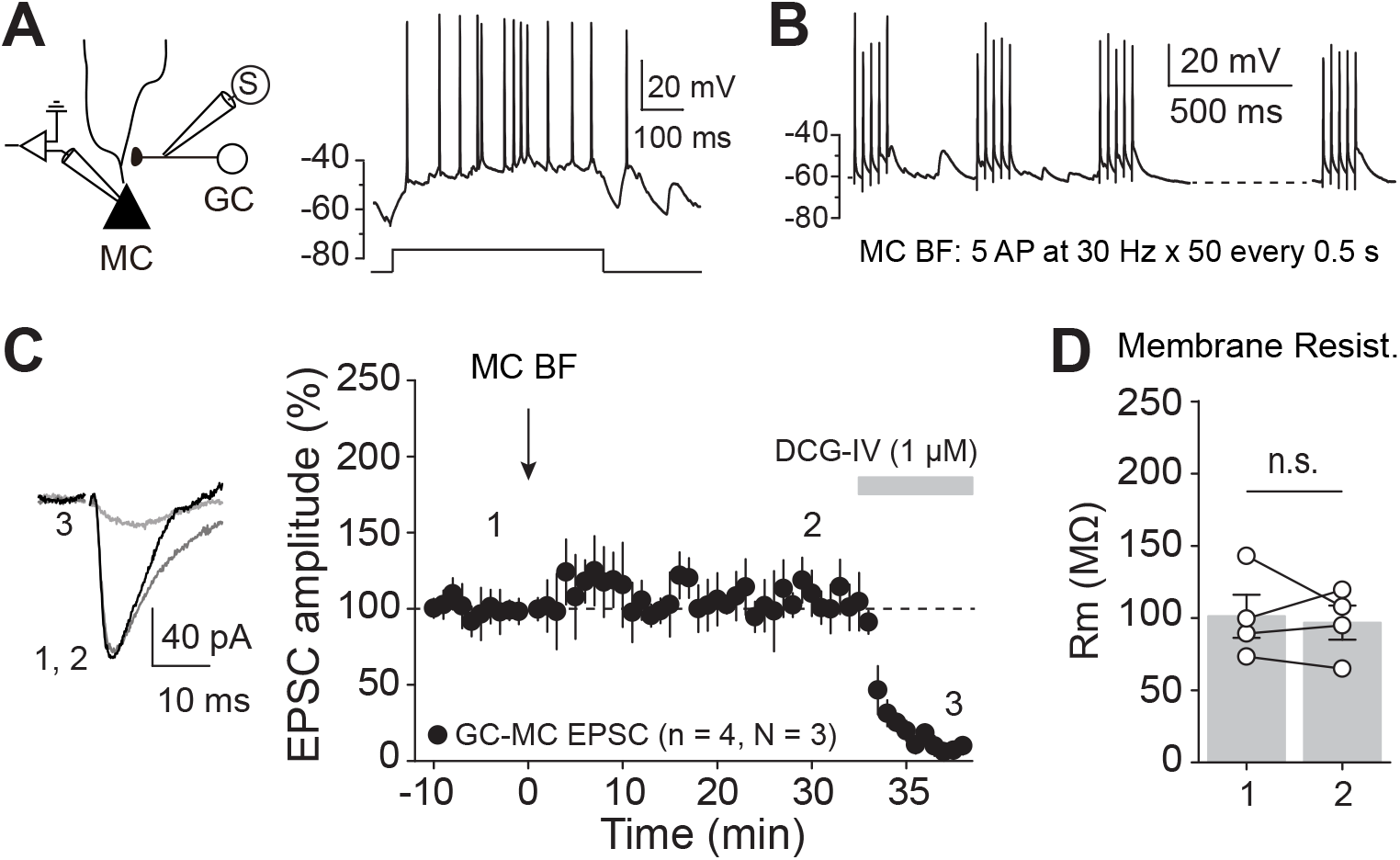
MC Burst Firing did not induce long-term changes at GC-MC synapse. **(A)** Left, diagram showing the recording conditions. Whole cell recordings were performed in MCs in response to GC axon stimulation, while both excitatiory and inhibitory transmission were left intact. Right, voltage response elicited by a depolarizing current step injection (shown below) illustrating typical MC firing pattern. **(B)** Current clamp recordings showing the induction protocol. MC Burst Firing (MC BF) was composed of 5 AP at 30 Hz, repeated 50 times, every 0.5 s. **(C)** Representative traces and time course summary plot showing that application of MC BF did not induce any significant long-term change of GC-MC EPSC amplitude (105.9 ± 10.7 % of baseline, n = 4, p > 0.05, paired t-test). DCG-IV was applied at the end of each experiment to confirm GC input nature. **(D)** MC membrane resistance was not affected by MC BF application (before MC BS: 101.3 ± 14.9 MΩ, after MC BS: 96.8 ± 11.8 MΩ, before *vs* after MC BS: p > 0.05, paired t-test, n = 4).

## Notes

### Competing Interest Statement

The authors have declared no competing interest.

